# High activity of an affinity-matured ACE2 decoy against Omicron SARS-CoV-2 and pre-emergent coronaviruses

**DOI:** 10.1101/2022.01.17.476672

**Authors:** Joshua J. Sims, Sharon Lian, Rosemary L. Meggersee, Aradhana Kasimsetty, James M. Wilson

## Abstract

The viral genome of severe acute respiratory syndrome coronavirus 2 (SARS-CoV-2), particularly its cell-binding spike protein gene, has undergone rapid evolution during the coronavirus disease 2019 (COVID-19) pandemic. Variants including Omicron BA.1 and Omicron BA.2 now seriously threaten the efficacy of therapeutic monoclonal antibodies and vaccines that target the spike protein. Viral evolution over a much longer timescale has generated a wide range of genetically distinct *sarbecoviruses* in animal populations, including the pandemic viruses SARS-CoV-2 and SARS-CoV-1. The genetic diversity and widespread zoonotic potential of this group complicates current attempts to develop drugs in preparation for the next *sarbecovirus* pandemic. Receptor-based decoy inhibitors can target a wide range of viral strains with a common receptor and may have intrinsic resistance to escape mutant generation and antigenic drift. We previously generated an affinity-matured decoy inhibitor based on the receptor target of the SARS-CoV-2 spike protein, angiotensin-converting enzyme 2 (ACE2), and deployed it in a recombinant adeno-associated virus vector (rAAV) for intranasal delivery and passive prophylaxis against COVID-19. Here, we demonstrate the exceptional binding and neutralizing potency of this ACE2 decoy against SARS-CoV-2 variants including Omicron BA.1 and Omicron BA.2. Tight decoy binding tracks with human ACE2 binding of viral spike receptor-binding domains across diverse clades of coronaviruses. Furthermore, in a coronavirus that cannot bind human ACE2, a variant that acquired human ACE2 binding was bound by the decoy with nanomolar affinity. Considering these results, we discuss a strategy of decoy-based treatment and passive protection to mitigate the ongoing COVID-19 pandemic and future airway virus threats.

**Author Summary:** Viral sequences can change dramatically during pandemics lasting multiple years. Likewise, evolution over centuries has generated genetically diverse virus families posing similar threats to humans. This variation presents a challenge to drug development, in both the breadth of achievable protection against related groups of viruses and the durability of therapeutic agents or vaccines during extended outbreaks. This phenomenon has played out dramatically during the coronavirus disease 2019 (COVID-19) pandemic. The highly divergent Omicron variants of severe acute respiratory syndrome coronavirus 2 (SARS-CoV-2) have upended previous gains won by vaccine and monoclonal antibody development. Moreover, ecological surveys have increasingly revealed a broad class of SARS-CoV-2-like viruses in animals, each poised to cause a future human pandemic. Here, we evaluate an alternative to antibody-based protection and prevention—a decoy molecule based on the SARS-CoV-2 receptor. Our engineered decoy has proven resistant to SARS-CoV-2 evolution during the ongoing COVID-19 pandemic and can neutralize all variants of concern, including Omicron BA.1 and Omicron BA.2. Furthermore, the decoy binds tightly to a broad class of *sarbecoviruses* related to pandemic SARS-CoV-2 and SARS-CoV-1, indicating that receptor decoys offer advantages over monoclonal antibodies and may be deployed during the COVID-19 pandemic and future coronavirus outbreaks to prevent and treat severe illness.

## 1. Introduction

Monoclonal antibody therapeutics with the ability to bind the spike protein of severe acute respiratory syndrome coronavirus 2 (SARS-CoV-2) and prevent cell entry have been critical tools in managing the coronavirus disease 2019 (COVID-19) pandemic (1, 2). These drugs prevent hospitalizations when applied early in the course of infection (3) and can provide critical passive protection for vulnerable populations of immunocompromised patients who cannot mount a protective response to vaccines (4). However, monoclonals have proven susceptible to SARS-CoV-2 evolution (5). This susceptibility may arise because the spike epitopes most sensitive to neutralization have been under intense selection as the virus has made gains in transmissibility and its ability to evade human immunity (6). Detailed structural analyses have highlighted that the SARS-CoV-2 receptor-binding domain (RBD) is obscured from antibodies by a thick glycan coat that it is particularly evident when the RBD is in the downward conformation, enabling it to evade neutralization. Additionally, the mobility of the RBD between up and down conformations may itself present an issue for RBD:antibody interactions (7). Thus, monoclonal antibodies that target the RBD in the upward conformation may show reduced susceptibility (7, 8).

Several groups have identified several sites present in the functional binding epitope of the RBD spike protein that have undergone mutations to evade all currently available classes of monoclonal antibodies (9-11). The most significant escape mutants identified are K417 for class 1 antibodies, L452R and E484 for class 2 antibodies, and R346, K444, and G446-450 sites for class 3 antibodies (9, 11). It is concerning that many of these mutations are present in emergent CoV-2 variants, which also show reduced susceptibility to monoclonal antibodies (9, 12, 13). While comprehensive structural analysis suggests potential improvements for targeting monoclonal antibodies and other biologics to the spike protein for inhibition (14-17), the fact remains that SARS-CoV-2 evolution during the course of the pandemic has generated variants which have evaded essentially the whole spectrum of clinical monoclonal candidates to date (5, 18). Furthermore, evidence suggests that application of single monoclonal antibody types in a therapeutic setting can rapidly give rise to escape mutants (19-22). Together, these findings call into question the ability of the antibody platform to keep pace with the course of the COVID-19 pandemic, or to be of use in future pandemics caused by other coronaviruses.

Receptor decoys may represent a mode of viral neutralization that is more resistant to continued viral evolution and escape-mutant generation (23). SARS-CoV-2 evolution has occurred in a way that retains tight binding to its primary cell entry receptor, angiotensin-converting enzyme 2 (ACE2) (24). We and others have developed affinity-matured, soluble ACE2 decoy molecules that potently neutralize SARS-CoV-2 (23, 25-30). Our soluble Fc-fused decoy, CDY14HL-Fc4, contains six amino acid substitutions that improve the neutralization of CoV-2 variants by 300-fold versus un-engineered ACE2 and an active site mutation that ablates its endogenous angiotensin-cleaving activity. Furthermore, CDY14HL maintains tight binding or neutralizing activity for the distantly related betacoronaviruses (*sarbecoviruses*) WIV1-CoV, and SARS-CoV-1 despite being engineered for improved activity against SARS-CoV-2 (25). This property suggests that this decoy may be a useful tool to combat future pandemics from currently pre-emergent, ACE2-dependent coronaviruses.

Here, we evaluate the binding and neutralization activity of CDY14HL against a wide range of emerging SARS-CoV-2 variants, including Omicron BA.1 and Omicron BA.2, and a wide range of pre-emergent coronaviruses with similarity to SARS-CoV-1 and SARS-CoV-2. These studies suggest the broad utility of decoy-based viral entry inhibitors in combating current and future coronavirus pandemics.

## 2. Results

### 2.1 CDY14HL maintains tight binding to diverse SARS-CoV-2 variants

We set out to evaluate the ability of our engineered ACE2 decoy to neutralize emerging SARS-CoV-2 strains. We previously assessed binding between the decoy and CoV RBDs by first expressing and purifying the RBDs and subjecting these to binding analysis using surface plasmon resonance (25). We sought an RBD binding assay that would allow us to assess a broad range of viral sequences for decoy compatibility quickly, without the need to express and purify unique viral proteins. As a first step, we assessed binding to variant RBDs using a yeast display system (31) (**Figure 1A**). We generated budding yeast displaying viral RBDs as fusion proteins to the cell-surface-tethered yeast protein Aga2. We then incubated the RBD yeast with CDY14HL-Fc1 fusion protein and assessed decoy binding via flow cytometry by staining bound decoy with a fluorescent secondary antibody. CDY14HL-Fc1 bound the ancestral (Wuhan-Hu-1) RBD with an apparent affinity of 0.14 nM (**Figure 1B and Table 1**). This result is in good agreement with the 0.21 nM affinity obtained for the interaction using an orthogonal technique, bio-layer interferometry (BLI, **Figure S1**), as well as with the picomolar binding affinity we previously measured for CDY14HL-Fc4:RBD interaction using surface plasmon resonance (25). Since our first description of CDY14HL (25), several SARS-CoV-2 variants of concern (VoCs) have emerged, some with greater transmissibility and clinical sequelae than the original Wuhan strain (13, 32, 33); most of the RBD evolution has occurred at the ACE2 interface (**Figures 1C and 1D**) and could impact decoy affinity. We used the yeast display system to evaluate decoy binding to RBDs from five of these VoCs (Iota, Delta, Delta Plus, Lambda, and Mu). CDY14HL maintained sub-nanomolar binding affinity for all VoC RBDs (**Figure 1B and Table 1**). This finding is consistent with the broad resistance of CDY14HL to SARS-CoV-2 variant evolution previously observed in binding and pseudotype neutralization studies (25).

**Table 1:** RBD binding constants and neutralization IC50 values for CDY14HL-Fc1 vs SARS-CoV-2 variants

| Strain | RBD mutations relative to Wuhan-Hu1 | Yeast assay RBD binding $K_D$ (nM) | Pseudotype neutralization assay IC50 (ng/ml) |
| --- | --- | --- | --- |
| CoV-2 (Wuhan-Hu-1) | none | 0.14 | 35 |
| Delta | L452R T478K | 0.21 | 14 |
| Delta Plus | K417N L452R T478K | 0.21 | 10 |
| Delta with 439K/484K/501Y | K417N N439K L452R T478K E484K N501Y | 0.07 | NC |
| Iota | E484K | 0.10 | NC |
| Kappa | L452R E484Q | NC | 18 |
| Lambda | L452Q F490S | 0.20 | 11 |
| Mu | R346K E484K N501Y | 0.10 | 21 |
| Omicron BA.1 | G339D S371L S373P S375F K417N N440K G446S S477N T478K E484A Q493K G496S Q498R N501Y Y505H | 0.031 | 18 |
| Omicron BA.2 | G339D S371F S373P S375F T376A D405N R408S K417N N440K S477N T478K E484A Q493R Q498R N501Y Y505H | 0.024 | 30 |
| Zeta | E484K | NC | 37 |

NC: Not Collected

**Fig 1.**
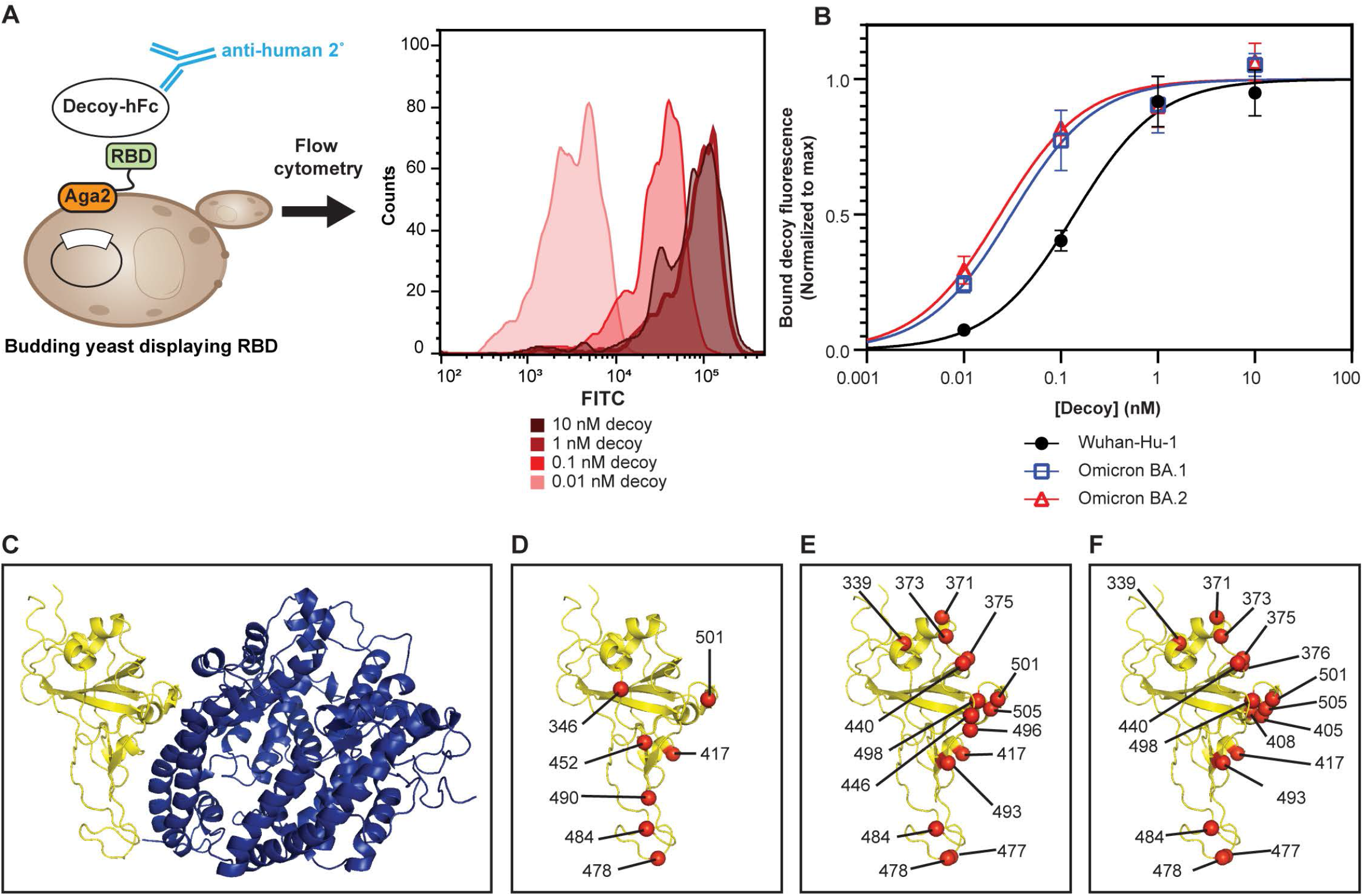
CDY14HL maintains tight binding to diverse SARS-CoV-2 variants. (A) Scheme for measuring decoy binding to yeast surface-expressed RBDs. Histograms are from a representative replicate of decoy binding to yeast-displayed Wuhan-Hu-1 RBD. (B) Representative decoy binding data for the RBD from the ancestral (Wuhan-Hu-1) SARS-CoV-2 strain and both Omicron variants. (C) The complex (68) between the SARS-CoV-2 Wuhan-Hu-1 RBD (yellow ribbons) and human ACE2 (blue ribbons). (D) The Wuhan-Hu-1 RBD from the ACE2 complex structure is shown alone. Red spheres indicate the α-carbon of amino acid residues mutated in several pre-Omicron SARS-CoV-2 variant strains. Amino acid position numbers are show for each mutated residue. These mutations cluster around the interface with ACE2. Wuhan-Hu-1 RBD is shown with the positions mutated in Omicron BA.1 (E) and Omicron.BA.2 (F) in red spheres.Error bars represent standard deviation of at least four replicate measurements.

Next, we examined the Omicron VoCs. Unlike previous variants, which contain one, two, or three RBD mutations, the Omicron BA.1 RBD differs from the ancestral strain by 15 amino acids (**Figure 1E**) (34), while the Omicron BA.2 variant RBD differs by 16 amino acids from the ancestral strain (**Figure 1F**). This level of mutation has caused a reduction in the efficacy of first-generation vaccines and most of the monoclonal antibodies developed for therapeutic and passive prophylaxis applications (35-40). Since our decoy was engineered for optimal binding to the ancestral strain, we sought to determine if this degree of antigenic drift would impact binding to the Omicron strains. Remarkably, the CDY14HL-Fc1 bound both Omicron strain RBDs with sub-nanomolar affinity as measured by the RBD yeast display assay (**Figure 1B** and **Table 1**). The decoy bound yeast-displayed Omicron RBDs at least four-fold more tightly than the ancestral RBD (0.03 nM and 0.02 nM for the BA.1 and BA.2 strains, respectively, **Table 1**). Another group working on their own decoy recently tested CDY14HL-Fc1 using a similar binding strategy, using HEK293 cells expressing Omicron BA.1 and BA.2 spike glycoprotein instead of yeast. Their results corroborate our data and highlighted that our decoy bound Omicron BA.1 and BA.2 at low nanomolar concentrations (18). We have also confirmed sub-nanomolar decoy affinity for the Omicron strain RBDs using BLI, though the two distinct assay formats produced different rank-orders of variant affinities (**Figure S1**).

### 2.2 CDY14HL maintains potent neutralization for diverse SARS-CoV-2 variants

We next investigated whether the broad decoy affinity for SARS-CoV-2 variants observed using the yeast display binding assay would translate to potent viral neutralization. We used lentiviruses harboring a luciferase reporter gene and pseudotyped with spike proteins from various SARS-CoV-2 strains to measure the neutralization potency (half-maximal inhibitory concentration [IC50]) of purified CDY14HL-Fc1 decoy. CDY14HL-Fc1 neutralized Omicron BA.1 and BA.2 more potently than the ancestral strain (18 ng/ml for BA.1 and 30 ng/ml for BA.2 vs. 35 ng/ml for Wuhan-Hu-1, **Figure 2 and Table 1**). We extended this approach to include 6 additional variants (Delta, Delta Plus, Kappa, Lambda, Mu, and Zeta). CDY14HL neutralized all SARS-CoV-2 strain pseudotypes tested with IC50 values near or below the potency of the ancestral strain, Wuhan, against which it was engineered (**Figure 2**). Together with these binding data, these variant neutralization data indicate that the CDY14HL decoy has a broad tolerance for CoV-2 strain evolution, losing virtually no potency in the face of extensive viral spike remodeling under the evolutionary pressures of a several-year-long global pandemic.

**Fig 2.**
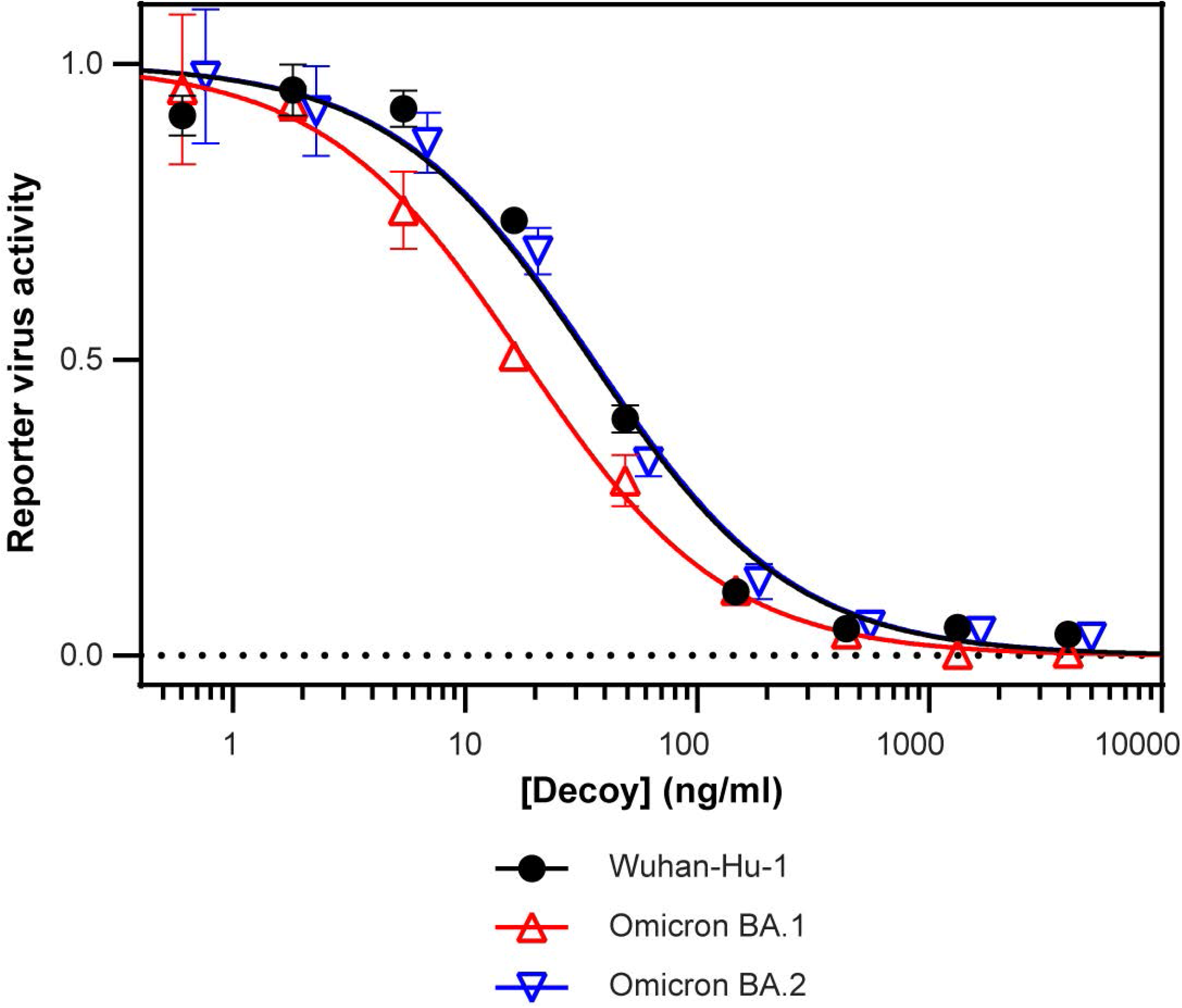
CDY14HL maintains potent neutralization for diverse SARS-CoV-2 variants. Viral neutralization assay using lentiviruses pseudotyped with the ancestral (Wuhan-Hu-1), Omicron BA.1, or Omicron BA.2 variant spike protein. Error bars represent standard deviation of at least three replicate measurements.

### 2.3 CDY14HL binds diverse ACE2-dependent CoVs

Given the broad activity towards SARS-CoV-2 variants with diverse spike sequences, we next evaluated the ability of CDY14HL to bind RBDs from a diverse set of coronaviruses with pandemic potential. Aside from SARS-CoV-1 and SARS-CoV-2, we identified 23 *sarbecoviruses* isolated from bats across Asia, Africa, and Europe (24, 41-53), many of which are thought to use ACE2 as a receptor (**Figure 3A**). We cloned synthetic RBD genes into the yeast display format for binding analysis (**Table 2**). Additionally, we included the RBD from the human coronavirus NL63, an alpha-CoV with a genetically distinct RBD that has been shown to use ACE2 for cell entry (54). We determined the binding affinities of yeast-displayed RBD to CDY14HL-Fc1 by flow cytometry.

**Table 2:**
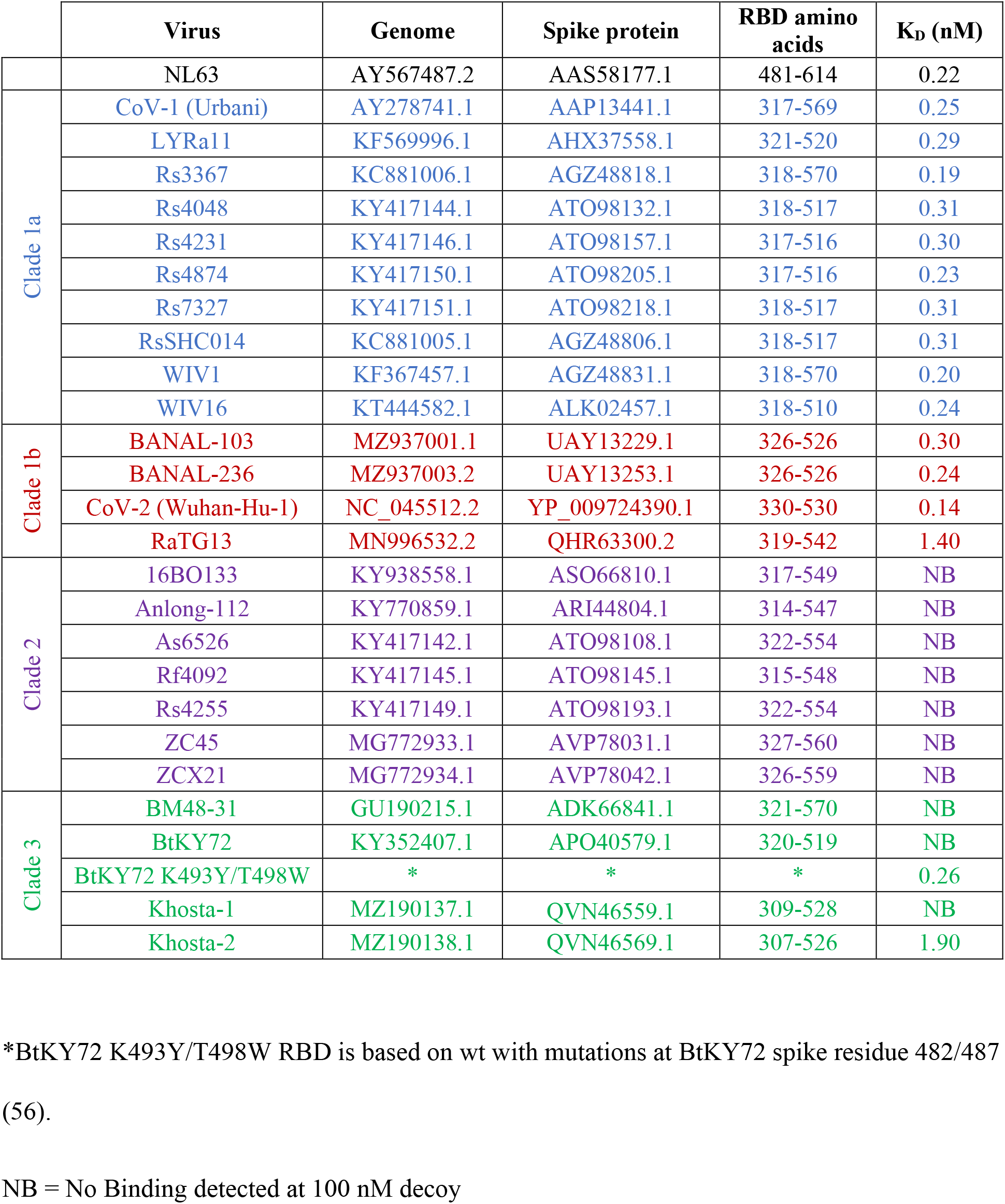
*Sarbecovirus* RBD sequence and decoy binding data

**Fig 3.**
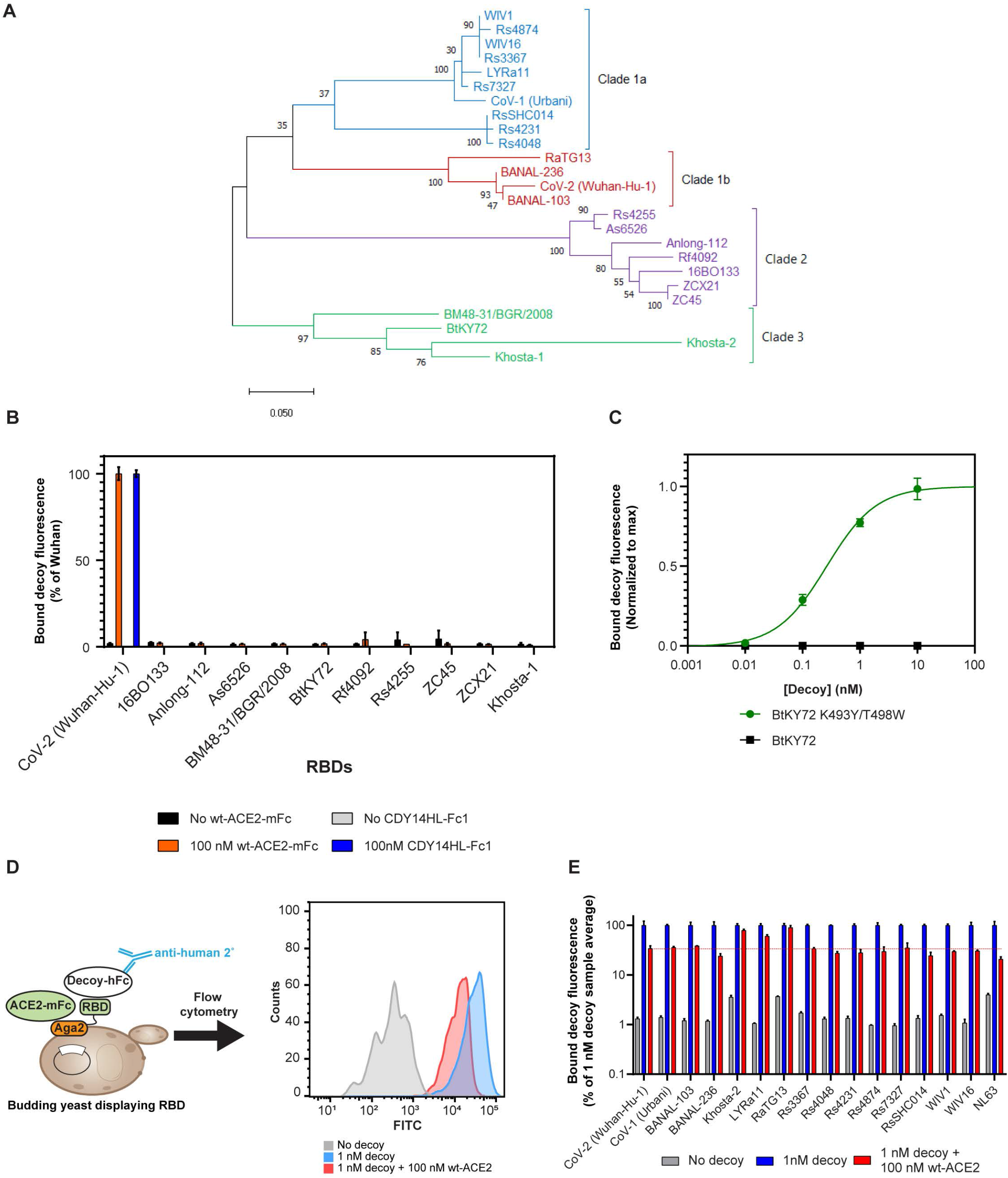
CDY14HL binds diverse ACE2-dependent CoVs. (A) Phylogenetic tree of *sarbecovirus* RBDs created using the maximum likelihood method. (B) Yeast displayed RBDs were used to test for decoy or wt-ACE2 binding at 100 nM. (C) Yeast displayed BtKY72 RBDs were titrated with decoy and evaluated to determine the dissociation equilibrium constant. (D) Schematic for measuring the competition between decoy and endogenous ACE2 receptor using yeast-displayed RBDs. Histogram data are from a representative replicate for the SARS-CoV-2 Wuhan-Hu-1 RBD. (E) Relative levels of decoy binding to diverse RBDs under several conditions, as assessed via the yeast display system. Error bars represent standard deviation of at least three replicate measurements.

Several patterns in decoy binding emerged. *Sarbecoviruses* from clades 1a and 1b (CoV-1-like and CoV-2-like CoVs, respectively, isolated from southern China and Laos) bound CDY14HL with sub-nanomolar affinities in the yeast display assay. The exception to this trend was RaTG13, which had a dissociation equilibrium constant of 1.4 nM (**Table 2**). Research has recently shown that RaTG13 has weaker binding to human ACE2 than other members of the clade (55). This behavior suggests that affinity for the decoy and the endogenous human ACE2 receptor are closely linked, as we have previously observed (25).

Except for Khosta-2 (56), *sarbecoviruses* from clades 2 and 3 (isolated in Asia and Europe/Africa, respectively) did not bind CDY14HL-Fc1. Based on a recent survey of RBD:ACE2 usage (56), we suspected these clade 2 and 3 RBDs were not capable of human ACE2 binding, which we confirmed in the yeast display system using a soluble wild-type human ACE2-Fc1 fusion protein at 100 nM (**Figure 3B**). Interestingly, Bloom and colleagues recently discovered that a K493Y/T498W double mutation to the RBD of the clade 3 CoV BtKY72 confers human ACE2 binding (56). We found that our decoy bound BtKY72 K493Y/T498W RBD with sub-nanomolar affinity, though it could not bind wt-BtKY72 (**Table 2, Figure 3C**). This suggests that if distant CoVs acquire mutations that confer the potential for zoonotic spread, CDY14HL would retain potent inhibitory potential.

These binding data are in broad agreement with our previous work demonstrating tight decoy binding to RBDs from SARS-CoV-1 and WIV1-CoV (25); indeed, we confirm SARS-CoV-1 and WIV1-CoV binding here using the yeast display assay (**Table 2**). However, we reasoned that viral neutralization comes about not only due to the decoy affinity for the viral target, but also from the ability of the decoy to compete for viral binding with the endogenous ACE2 receptor. To assess this possibility, we employed a competitive binding assay between the decoy and ACE2 receptor in the yeast system. We incubated RBD yeast with a low concentration of decoy (1 nM of CDY14HL-hFc1; 95 ng/ml) along with a 100-fold molar excess of wt-ACE2 (100 nM of wt-ACE2-mFc, with a mouse Fc fusion to distinguish it from the decoy). We assessed the level of decoy binding retained in the presence of receptor competition by flow cytometry and compared these values across the set of RBDs (**Figure 3D**).

The positive control, the RBD from the well-neutralized ancestral SARS-CoV-2 strain, retained 34% of decoy binding in the presence of 100-fold molar excess of wt-ACE2-mFc (**Figure 3E**). Similar to SARS-CoV-2, all clade 1a and 1b *sarbecovirus* RBDs tested retained at least 24% binding in the competition assay. Khosta-2, a clade 3 RBD with weak decoy binding (**Table 2**), retained 79% percent decoy binding in the presence of 100-fold excess wt-ACE2. Similarly, the relatively weak-binding RBD from RaTG13 also retained almost complete decoy binding under competition. These examples highlight that decoy affinity alone may be a poor determinant of the ability to compete with endogenous ACE2 as an inhibitor (**Figure S2**). The lone alpha-CoV in our study, NL63, retained the lowest fraction of decoy binding in the competition assay (21%, **Figure 3E**). Further study is needed to determine whether this result indicates a lower neutralizing potency of the decoy for the genetically distinct ACE2-dependent alpha-CoVs. Together with the observed sub-nanomolar decoy binding affinity, these competitive binding data predict broad and potent neutralization of beta-CoVs that can bind human ACE2. As many factors including subtle shifts in co-receptor usage and cell entry mechanism can define the potency of an inhibitor (57), further studies with diverse CoVs in pseudotyped and live-virus systems will be required to confirm this finding.

## 3. Discussion

Currently available COVID-19 treatments such as monoclonal antibodies are fast becoming less effective as new variants emerge (12, 58). Alternative protein-based viral neutralizers exist, including camelid-derived single chain antibodies. While these have several advantages over conventional antibodies with respect to small size, delivery modalities, and manufacturability (59-61), nanobodies still recognize a specific epitope on the SARS-CoV-2 RBD spike surface and thus are still susceptible to mutations that could eliminate this binding epitope.

We previously reported the development of an affinity-matured, soluble ACE2 decoy, termed CDY14HL. This decoy binds and neutralizes SARS-CoV-2 strains from the early pandemic as well as the related pandemic *sarbecovirus*, SARS-CoV-1 (25). In this study, we have shown that the affinity-matured decoy retains broad neutralizing activity against every SARS-CoV-2 variant tested, including Delta, Delta Plus, Omicron BA.1, and Omicron BA.2. These results highlight a major advantage of receptor-based decoy molecules over other viral spike inhibitor biologics; unlike monoclonal antibodies, decoy binding of a variant is tightly linked to receptor binding and thus viral fitness through evolution. Zhang *et al*., illustrated this very effectively through their own affinity experiments where they compared engineered decoy binding of omicron BA.1 and BA.2 to four monoclonal antibodies authorized for clinical use (18). They determined that REGN10933 and LY-CoV555 had no binding activity to either Omicron BA.1 or BA.2. REGN10987 was found to bind to Omicron BA.2 weakly and Omicron BA.1 not at all. However, the most surprising result was for VIR-7831, a monoclonal targeting a distinct and highly conserved spike site that was selected to yield both broad sarbecovirus neutralization and resistance to SARS-CoV-2 variant evolution (62). VIR-7831 was only able to bind to Omicron BA.1 and very weakly to BA.2, a fact that subsequently led to the removal of FDA authorization for the drug (63).

Our original strategy for deploying the decoy was developed in the context of preventing SARS-CoV-2 infection. We accomplished this aim through the creation of a recombinant adeno-associated virus (rAAV) vector expressing the decoy that is administered nasally to engineer proximal airway cells to express neutralizing levels of the decoy at the airway surface (i.e., the virus’s entry point). This approach could be particularly useful for immunocompromised patients who do not generate protective immunity following active vaccination. We are also developing the decoy as a therapeutic protein for treatment, or possibly prevention in high-risk groups, following parenteral administration.

The relentless emergence of new, highly transmissible SARS-CoV-2 variants in the current pandemic reminds us of our vulnerability to the power of zoonosis and the intense selection pressures experienced by pandemic viruses to evolve into more pathogenic and/or transmissible variants. This experience suggests the importance of proactively developing countermeasures against future pandemics, which will likely be caused by a coronavirus based on the recent history of SARS, Middle East respiratory syndrome, and COVID-19. Indeed, CoVs constitute a major fraction of pre-zoonotic viruses ranked by multiple genetic and environmental factors for pandemic potential (64). This threat compelled us to evaluate the competitive binding of our ACE2 decoy to spike proteins from a variety of animal coronaviruses with zoonotic potential, particularly *sarbecoviruses* that can bind human ACE2, or that whose viral spike proteins only need a small number of mutations to acquire human ACE2 binding. We were delighted to find that the decoy retained very high binding activity against spike proteins from every pre-emergent strain studied that also had the ability to bind human ACE2, suggesting the broad utility of the decoy in the current and future coronavirus pandemics.

COVID-19 has illustrated how powerful the drive for viral fitness can be in circumventing immunity generated from previous infection, vaccines, and antibody therapeutics. This rapid evolution is substantially amplified in the setting of a global pandemic caused by a highly transmissible virus. The use of a decoy protein based on a soluble version of a viral receptor holds the promise of significantly restricting viral escape, as any mutation that diminishes decoy binding will likely also diminish receptor binding and thus viral fitness. We are quickly moving this ACE2 decoy into the clinic in the AAV platform as well as a protein therapeutic as a possible solution to COVID-19 variants and to prepare for future coronavirus outbreaks.

## 4. Materials and Methods

### 4.1 CoV pseudotyped lentiviral neutralization assay

Replication-incompetent lentiviruses pseudotyped with full-length CoV spike proteins and packaging for a Renilla luciferase reporter gene were purchased from Integral Molecular: RVP-701L Wuhan (lot CL-114B), RVP-763L Delta (lot CL-267A), RVP-736L Zeta (lot CL-255A), RVP-730L Kappa (lot CL-247A), RVP-768L Omicron BA.1 (lot CL-297A), RVP-770L Omicron BA.2 (lot CL-310), RVP-767L Mu (lot CL-274A), RVP-766L Lambda (lot CL-259A), and RVP-765L Delta Plus (lot CL-258A). They have been shown to be suitable for neutralization measurements with a high linear correlation of IC50s measured by the pseudotyped virus neutralization assay with SARS-CoV-2 PRNT (65). We performed neutralization assays using human embryonic kidney 293T cells overexpressing ACE2 (Integral Molecular) as previously described (25), performing a minimum of 3 replicates to arrive at the data presented.

### 4.2 Recombinant protein production

To generate wt-ACE2-mFc for competitive binding assays, we cloned human ACE2 (1–615) fused to a C-terminal mouse IgG2a Fc into pcDNA3.1. We transfected the plasmid into Expi293 cells for expression. The supernatant was collected and exchanged to 0.1 M sodium phosphate, pH 7.2 and 150 mM NaCl buffer for purification on Protein A Sepharose 4B (ThermoFisher). The protein was eluted in 0.1 M citric acid, pH 3.0 and neutralized in 1 M Tris, pH 9.0 before a final buffer exchange to 25 mM HEPES pH 7.2 and 150 mM NaCl by size-exclusion chromatography with Superose 6 resin (Cytiva). For these studies, we cloned the engineered CDY14HL 1–615 fragment in front of the human IgG1 Fc domain for expression and purification. We previously characterized a decoy fusion to human IgG4 Fc (25), but found that Fc1 and Fc4 decoy fusions behave similarly with respect to binding and neutralization (e.g., the IC50 values against Wuhan-Hu1 pseudotypes were 37 ng/ml and 35 ng/ml for CDY14HL-Fc4 and CDY14HL-Fc1, respectively).

### 4.3 Phylogenic tree construction

The RBD sequences of the CoVs were taken from spike protein coding sequences (**Table 2**) downloaded from the National Center for Biotechnology Information (NCBI). Using MEGA X (66), we aligned the amino acid sequences in ClustalW and constructed a phylogenic tree using maximum likelihood analysis and bootstrapping with 100 replicates.

### 4.4 Yeast display binding assays

The nucleic acid sequences of the CoV RBDs were taken from NCBI. Accession numbers and domain boundary details are included in **Table 2**.We cloned the RBDs into a plasmid between an upstream yeast Aga2 gene and a downstream hemagglutinin (HA) epitope tag with flexible GSG linkers. The plasmid has a low-copy centromeric origin similar to that of pTCON2 (67). Plasmids were transformed into EBY100 using the Frozen-EZ Yeast Transformation II Kit (Zymo). We grew colonies in SD-Trp media before induction in log phase for 24 hr at 30°C in SG-CAA (67). To determine K_D_, yeast were incubated with a 1:10 dilution series of CDY14HL-Fc1 for 16 hr at 25°C. To check for ACE2 binding, yeast were incubated with 100 nM of CDY14HL-Fc1 or wt-ACE2-mFc for 16 hr at 25°C. For the competition binding, yeast were incubated with 1 nM of CDY14HL-Fc1 and 100 nM of wt-ACE2-mFc for 30 min at 25°C. Afterwards, samples were washed and stained for flow cytometry using appropriate seconday antibodies: goat anti-human fluorescein isothiocyanate (FITC; ThermoFisher A18812), anti-mouse Alexa Fluor 488 (Cell Signaling Technology 4408S), or rabbit anti-HA-PE (Cell Signaling Technology 14904S). We used phosphate-buffered saline with 0.1% bovine serum albumin for all staining and washes. For the titration of CDY14HL-Fc1, we incubated the yeast with 1:10 dilution series of CDY14HL-Fc1 at 25°C for 6 hr. The yeast were analyzed on an ACEA NovoCyte flow cytometer. We determined the level of CDY14HL-Fc1 binding by taking the mean FITC signal for 500 RBD+ yeast cells collected for each condition. We collected a minimum of 4 replicate binding curves to establish dissociation constants for each RBD variant. Since different RBDs expressed to slightly different levels on the surface of the yeast, for the purposes of plotting multiple binding curves in the same figure we normalized bound decoy fluorescence for each curve to its fit maximum value. We fitted the decoy concentration versus the decoy binding signal in GraphPad Prism using a three-parameter fit to the binding isotherm.

## Data Availability Statement

All relevant data are within the manuscript.

## Ethics Statement

N/A - IRB approval nor ethics approval were required as no medical samples, records, or animals were utilized in this study

## Acknowledgments

We thank Henry Hoff, Adam Salazar, Simin Zaidi, Kenneth Yancey, Dmitriy Fedorenko, Jasmine Tan, Yida He, Philip McGurk, and Sai Prasad Ganesh for protein expression and purification in support of these studies. We thank Alex Martino for assistance with BLI studies. We thank Nathan Denton for assistance with manuscript preparation and graphics.

## Author Contributions

J.J.S. **–** conceptualization, data curation, formal analysis, investigation, methodology, project administration, resources, supervision, validation, visualization, writing-original, writing-review, and edits. S.L. – conceptualization, data curation, formal analysis, investigation, methodology, resources, validation, visualization, writing-original, writing-review, and edits. R.M. data curation, formal analysis, investigation, project administration, visualization, writing-original, writing-review, and edits. A.K. formal analysis, investigation, visualization, writing-original, writing-review, and edits J.M.W. – conceptualization, funding acquisition, writing-original, writing-review, and edits.

## Supplemental Materials

**Figure S1.**
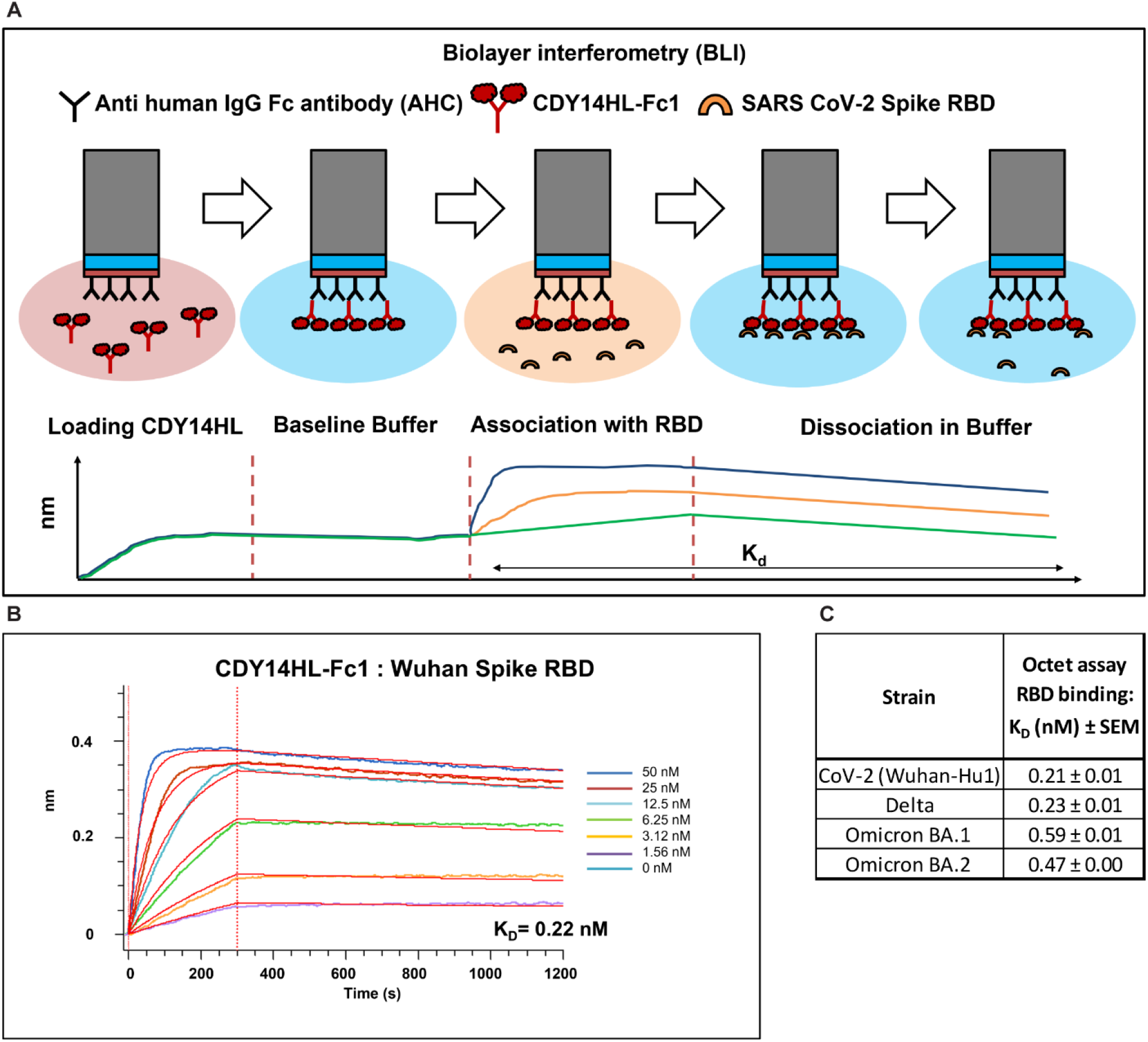
Biolayer interferometry (BLI) kinetics for CDY14HL-Fc1 and SARS-CoV-2 Variants. **A)** Depiction of Biolayer interferometry assay format. CDY14HL-Fc1 was immobilized as ligand and SARS-CoV-2 variant RBD was used as analyte **B)** Representative fitted sensogram for CDY14HL-Fc1 and CoV-2 Wuhan-Hu-1 spike RBD **C)** Table of K_D_ values (nM) for CDY14HL-Fc1 and four SARS-CoV-2 variant RBDs. Mean values and standard error of the mean (SEM) determined from three technical replicates.

**Figure S2.**
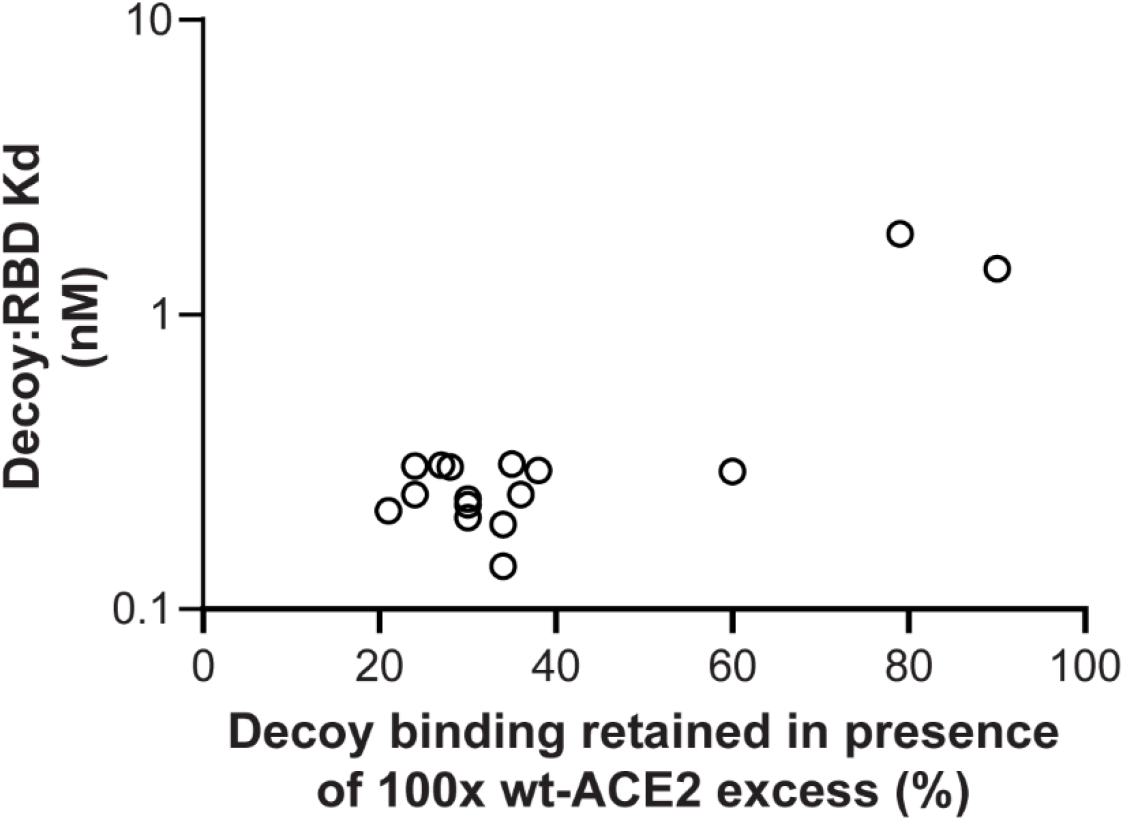
We use the data sets generated in **Table 2** and **Figure 3E** to evaluate the relationship between decoy affinity and the amount of decoy retained in a competitive binding assay.

## Materials and Methods

### Biolayer Interferometry

For the determination of binding affinity, biolayer interferometry measurements were taken on the Octet HTX (Sartorius Corporation). Purified CDY14HL-Fc1 was immobilized on an anti-human IgG Fc sensor (AHC sensor, Sartorius Corporation). Decoy loaded tips were dipped into wells containing purified His-tagged SARS-CoV-2 Spike RBD proteins (R&D Biotechne Corporation). RBD concentrations of 50, 25, 12.5, 6,25, 3.125 and 1.56 nM were used to determine the K_D_s. The diluent and running buffer for all experiments was 1x Kinetics Buffer (FortéBio, PBS+ 0.02% Tween20, 0.1% BSA, 0.05% sodium azide). K_D_ fits were determined by 1:1 binding model global grouped fits on the Octet Data Analysis HT Version 12.0.2.59 (FortéBio). All measurements were conducted in triplicate.

### Plasmid and sequence details for constructs used in the main text

Nucleotide sequence of CDY14HL-Fc1:

ATGTCAAGCTCTTCCTGGCTCCTTCTCAGCCTTGTTGCTGTAACTGCTGCTCAGTCCACCATTGAGGAACAGGCCAAGACATTTTTGGACAtGTTTAACCACaAAGCCGAAGACCTGTTCTATCAAAGTTCACTTGCTgCTTGGAATTATAACACCAATATTACTGAAGAGAATGTCCAAAACATGAATAAcGCTGGGGACAAATGGTCTGCCTTTTTAAAGGAACAGTCCACAtTTGCCCAAATGTATCCACTACAAGAAATTCAGAATCcCACAGTCAAGCTTCAGCTGCAGGCTCTTCAGCAAAATGGGTCTTCAGTGCTCTCAGAAGACAAGAGCAAACGGTTGAACACAATTCTAAATACAATGAGCACCATCTACAGTACTGGAAAAGTTTGTAACCCgGATAATCCACAAGAATGCTTATTACTTGAACCAGGTTTGAATGAAATAATGGCAAACAGTTTAGACTACAATGAGAGGCTCTGGGCTTGGGAAAGCTGGAGATCTGAGGTCGGCAAGCAGCTGAGGCCATTATATGAAGAGTATGTGGTCTTGAAAAATGAGATGGCgAGAGCAAAcCATTATGAGGACTATGGGGATTATTGGAGAGGAGACTATGAAGTAAATGGGGTAGATGGCTATGACTACAGCCGCGGCCAGTTGATTGAAGATGTGGAACATACCTTTGAAGAGATTAAACCATTATATGAACATCTTCATGCCTATGTGAGGGCAAAGTTGATGAATGCCTATCCTTCCTATATCAGTCCAATTGGATGCCTCCCTGCTCATTTGCTTGGTGATATGTGGGGTAGATTTTGGACAAATCTGTACTCTTTGACAGTTCCCTTcGGACAGAAACCAAACATAGATGTTACTGATGCAATGGTGGACCAGGCCTGGGATGCACAGAGAATATTCAAGGAGGCCGAGAAGTTCTTTGTATCTGTTGGTCTTCCTAATATGACTCAAGGATTCTGGGAAtATTCCATGCTAACGGACCCAGGAAATGTTCAGAAAGCAGTCTGCCtTCCCACAGCTTGGGACCTGGGGAAGGGCGACTTCAGGATCCTTATGTGCACAAAGGTGACAATGGACGACTTCCTGACAGCTCATCATGAGATGGGGCATATCCAGTATGATATGGCATATGCTGCACAACCTTTTCTGCTAAGAAATGGAGCTAATGAAGGATTCCATGAAGCTGTTGGGGAAATCATGTCACTTTCTGCAGCCACACCTAAGCATTTAAAATCCATTGGTCTTCTGTCACCCGATTTTCAAGAAGACAATGAAACAGAAATAAACTTCCTGCTCAAACAAGCACTCACGATTGTTGGGACTCTGCCATTTACTTACATGTTAGAGAAGTGGAGGTGGATGGTCTTTAAAGGGGAAATTCCCAAAGACCAGTGGATGAAAAAGTGGTGGGAGATGAAGCGAGAGATAGTTGGGGTGGTGGAACCTGTGCCCCATGATGAAACATACTGTGACCCCGCATCTCTGTTCCATGTTTCTAATGATTACTCATTCATTCGATATTACACAAGGACCCTTTACCAATTCCAGTTTCAAGAAGCACTTTGTCAAG CAGCTAAACATGAAGGCCCTCTGCACAAATGTGACATCTCAAACTCTACAGAAGCTGGACAGAAACTGTTCAATATGCTGAGGCTTGGAAAATCAGAACCCTGGACCCTAGCATTGGAAAATGTTGTAGGAGCAAAGAACATGAATGTAAGGCCACTGCTCAACTACTTTGAGCCCTTATTTACCTGGCTGAAAGACCAGAACAAGAATTCTTTTGTGGGATGGAGTACCGACTGGAGTCCATATGCAGACggaagtGACAAGACCCACACCTGTCCTCCATGTCCTGCTCCAGAACTGCTCGGCGGACCTTCCGTGTTCCTGTTTCCTCCAAAGCCTAAGGACACCCTGATGATCAGCAGAACCCCTGAAGTGACCTGCGTGGTGGTGGATGTGTCCCACGAGGATCCCGAAGTGAAGTTCAATTGGTACGTGGACGGCGTGGAAGTGCACAACGCCAAGACCAAGCCTAGAGAGGAACAGTACAACAGCACCTACAGAGTGGTGTCCGTGCTGACCGTGCTGCACCAGGATTGGCTGAACGGCAAAGAGTACAAGTGCAAGGTGTCCAACAAGGCCCTGCCTGCTCCTATCGAGAAAACCATCAGCAAGGCCAAGGGCCAGCCTAGGGAACCCCAGGTTTACACACTGCCTCCAAGCAGGGACGAGCTGACCAAGAATCAGGTGTCCCTGACCTGCCTCGTGAAGGGCTTCTACCCTTCCGATATCGCCGTGGAATGGGAGAGCAACGGCCAGCCTGAGAACAACTACAAGACAACCCCTCCTGTGCTGGACAGCGACGGCTCATTCTTCCTGTACAGCAAGCTGACAGTGGACAAGTCCAGGTGGCAGCAGGGCAACGTGTTCAGCTGCAGCGTGATGCACGAGGCCCTGCACAACCACTACACCCAGAAGTCCCTGAGCCTGTCTCCTGGAaaaTGATGA

Nucleotide sequence of wt-ACE2-mFc (human ACE2 1-615 fused to Fc from mouse IgG2A)

ATGTCAAGCTCTTCCTGGCTCCTTCTCAGCCTTGTTGCTGTAACTGCTGCTCAGTCCACCATTGAGGAACAGGCCAAGACATTTTTGGACAAGTTTAACCACGAAGCCGAAGACCTGTTCTATCAAAGTTCACTTGCTTCTTGGAATTATAACACCAATATTACTGAAGAGAATGTCCAAAACATGAATAATGCTGGGGACAAATGGTCTGCCTTTTTAAAGGAACAGTCCACACTTGCCCAAATGTATCCACTACAAGAAATTCAGAATCTCACAGTCAAGCTTCAGCTGCAGGCTCTTCAGCAAAATGGGTCTTCAGTGCTCTCAGAAGACAAGAGCAAACGGTTGAACACAATTCTAAATACAATGAGCACCATCTACAGTACTGGAAAAGTTTGTAACCCAGATAATCCACAAGAATGCTTATTACTTGAACCAGGTTTGAATGAAATAATGGCAAACAGTTTAGACTACAATGAGAGGCTCTGGGCTTGGGAAAGCTGGAGATCTGAGGTCGGCAAGCAGCTGAGGCCATTATATGAAGAGTATGTGGTCTTGAAAAATGAGATGGCAAGAGCAAATCATTATGAGGACTATGGGGATTATTGGAGAGGAGACTATGAAGTAAATGGGGTAGATGGCTATGACTACAGCCGCGGCCAGTTGATTGAAGATGTGGAACATACCTTTGAAGAGATTAAACCATTATATGAACATCTTCATGCCTATGTGAGGGCAAAGTTGATGAATGCCTATCCTTCCTATATCAGTCCAATTGGATGCCTCCCTGCTCATTTGCTTGGTGATATGTGGGGTAGATTTTGGACAAATCTGTACTCTTTGACAGTTCCCTTTGGACAGAAACCAAACATAGATGTTACTGATGCAATGGTGGACCAGGCCTGGGATGCACAGAGAATATTCAAGGAGGCCGAGAAGTTCTTTGTATCTGTTGGTCTTCCTAATATGACTCAAGGATTCTGGGAAAATTCCATGCTAACGGACCCAGGAAATGTTCAGAAAGCAGTCTGCCATCCCACAGCTTGGGACCTGGGGAAGGGCGACTTCAGGATCCTTATGTGCACAAAGGTGACAATGGACGACTTCCTGACAGCTCATCATGAGATGGGGCATATCCAGTATGATATGGCATATGCTGCACAACCTTTTCTGCTAAGAAATGGAGCTAATGAAGGATTCCATGAAGCTGTTGGGGAAATCATGTCACTTTCTGCAGCCACACCTAAGCATTTAAAATCCATTGGTCTTCTGTCACCCGATTTTCAAGAAGACAATGAAACAG AAATAAACTTCCTGCTCAAACAAGCACTCACGATTGTTGGGACTCTGCCATTTACTTACATGTTAGAGAAGTGGAGGTGGATGGTCTTTAAAGGGGAAATTCCCAAAGACCAGTGGATGAAAAAGTGGTGGGAGATGAAGCGAGAGATAGTTGGGGTGGTGGAACCTGTGCCCCATGATGAAACATACTGTGACCCCGCATCTCTGTTCCATGTTTCTAATGATTACTCATTCATTCGATATTACACAAGGACCCTTTACCAATTCCAGTTTCAAGAAGCACTTTGTCAAGCAGCTAAACATGAAGGCCCTCTGCACAAATGTGACATCTCAAACTCTACAGAAGCTGGACAGAAACTGTTCAATATGCTGAGGCTTGGAAAATCAGAACCCTGGACCCTAGCATTGGAAAATGTTGTAGGAGCAAAGAACATGAATGTAAGGCCACTGCTCAACTACTTTGAGCCCTTATTTACCTGGCTGAAAGACCAGAACAAGAATTCTTTTGTGGGATGGAGTACCGACTGGAGTCCATATGCAGACCCCAGAGGGCCCACAATCAAGCCCTGTCCTCCATGCAAATGCCCAGCACCTAACCTCTTGGGTGGACCATCCGTCTTCATCTTCCCTCCAAAGATCAAGGATGTACTCATGATCTCCCTGAGCCCCATAGTCACATGTGTGGTGGTGGATGTGAGCGAGGATGACCCAGATGTCCAGATCAGCTGGTTTGTGAACAACGTGGAAGTACACACAGCTCAGACACAAACCCATAGAGAGGATTACAACAGTACTCTCCGGGTGGTCAGTGCCCTCCCCATCCAGCACCAGGACTGGATGAGTGGCAAGGAGTTCAAATGCAAGGTCAACAACAAAGACCTCCCAGCGCCCATCGAGAGAACCATCTCAAAACCCAAAGGGTCAGTAAGAGCTCCACAGGTATATGTCTTGCCTCCACCAGAAGAAGAGATGACTAAGAAACAGGTCACTCTGACCTGCATGGTCACAGACTTCATGCCTGAAGACATTTACGTGGAGTGGACCAACAACGGGAAAACAGAGCTAAACTACAAGAACACTGAACCAGTCCTGGACTCTGATGGTTCTTACTTCATGTACAGCAAGCTGAGAGTGGAAAAGAAGAACTGGGTGGAAAGAAATAGCTACTCCTGTTCAGTGGTCCACGAGGGTCTGCACAATCACCACACGACTAAGAGCTTCTCCCGGACTCCGGGTAAATGA

### Map of typical Aga2-RBD-HA construct used for yeast display

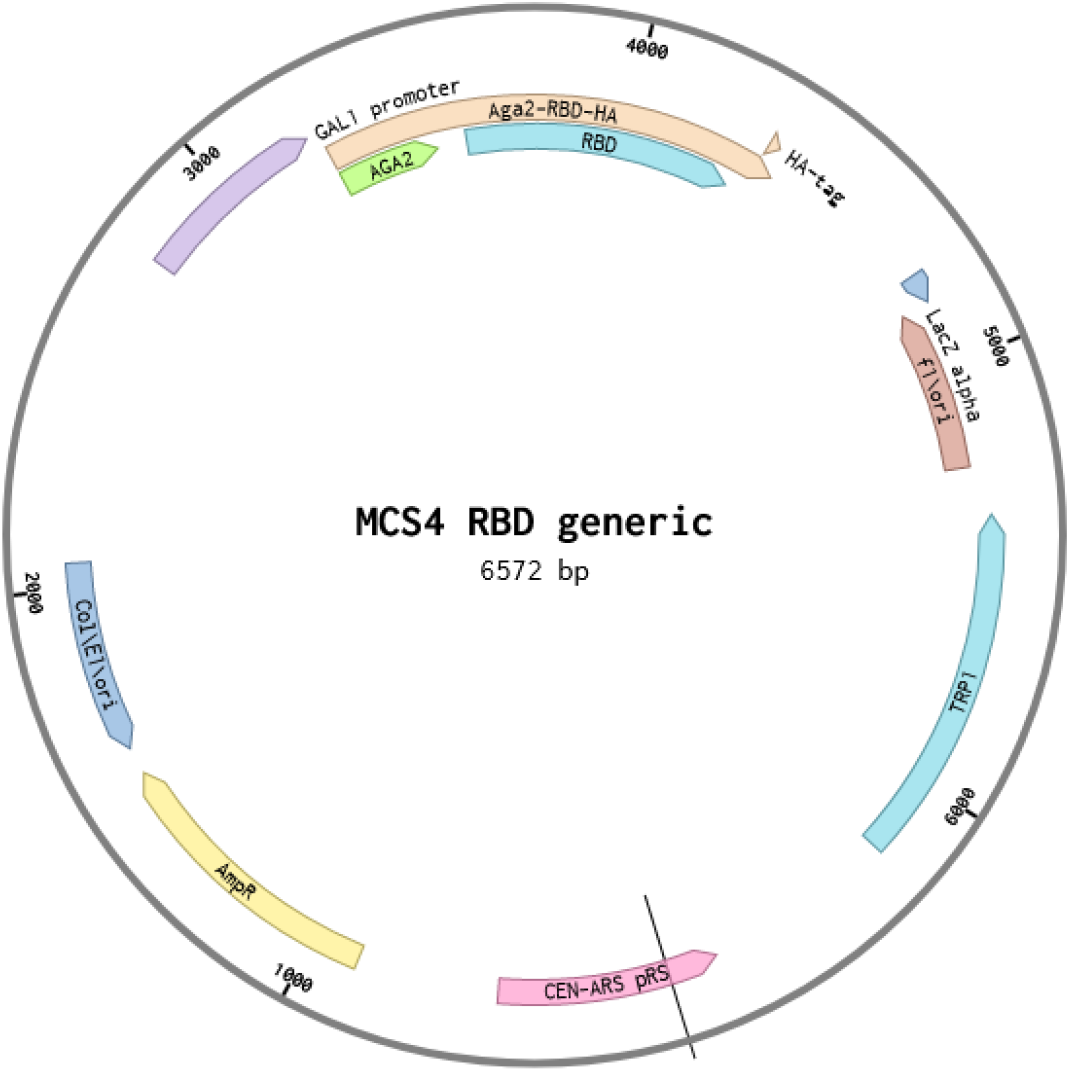

Sequence of Aga2-Wuhan-Hu-1-RBD-HA

ATGCAGTTACTTCGCTGTTTTTCAATATTTTCTGTTATTGCTTCAGTTTTAGCACAGGAACTGACAACTATATGCGAGCAAATCCCCTCACCAACTTTAGAATCGACGCCGTACTCTTTGTCAACGACTACTATTTTGGCCAACGGGAAGGCAATGCAAGGAGTTTTTGAATATTACAAATCAGTAACGTTTGTCAGTAATTGCGGTTCTCACCCCTCAACAACTAGCAAAGGCAGCCCCATAAACACACAGTATGTTTTTAAGGACAATAGCTCGACGATTGAAGGTAGAGGATCCGGAGGTAGCGGATCAGGAGGTGGAGGCTCCGGAGGAGGTGGACCAAACATCACCAATCTGTGCCCCTTCGGCGAAGTTTTCAACGCCACCAGATTCGCCAGCGTGTACGCCTGGAATAGAAAGCGGATCAGCAACTGTGTGGCCGACTACAGCGTGCTGTATAACAGCGCCAGCTTCAGCACATTCAAGTGCTACGGCGTGTCCCCTACCAAGCTGAACGACCTGTGTTTTACCAACGTGTACGCCGATAGCTTCGTGATTAGAGGCGACGAGGTGCGGCAGATCGCCCCTGGCCAGACCGGTAAGATCGCTGATTACAACTACAAGCTGCCTGACGACTTCACCGGCTGTGTGATCGCCTGGAACAGCAACAACCTGGACAGCAAGGTGGGCGGCAACTATAACTACCTGTACCGGCTGTTCAGAAAATCTAATCTGAAGCCCTTCGAGAGAGATATCAGCACAGAGATCTACCAGGCCGGCAGCACCCCTTGCAACGGCGTGGAAGGCTTTAACTGCTACTTCCCCCTGCAGAGCTACGGATTCCAGCCAACAAATGGAGTCGGCTACCAACCTTACAGAGTGGTCGTGCTCTCCTTTGAGCTGCTGCACGCCCCTGCCACAGTGTGCGGACCTAAGAAAAGCGGATCTTCTGGATCTGGATCTGGATCTGAATCTAAGTCTACTGGATACCCATACGACGTTCCAGACTACGCTCTGCAG GCTAGTGGTTGA

